# Gemini: Memory-efficient integration of hundreds of gene networks with high-order pooling

**DOI:** 10.1101/2023.01.21.525026

**Authors:** Addie Woicik, Mingxin Zhang, Hanwen Xu, Sara Mostafavi, Sheng Wang

## Abstract

**Motivation:** The exponential growth of genomic sequencing data has created ever-expanding repositories of gene networks. Unsupervised network integration methods are critical to learn informative representations for each gene, which are later used as features for downstream applications. However, these network integration methods must be *scalable* to account for the increasing number of networks and *robust* to an uneven distribution of network types within hundreds of gene networks.

**Results:** To address these needs, we present Gemini, a novel network integration method that uses memory-efficient high-order pooling to represent and weight each network according to its uniqueness. Gemini then mitigates the uneven distribution through mixing up existing networks to create many new networks. We find that Gemini leads to more than a 10% improvement in F_1_ score, 14% improvement in micro-AUPRC, and 71% improvement in macro-AURPC for protein function prediction by integrating hundreds of networks from BioGRID, and that Gemini’s performance significantly improves when more networks are added to the input network collection, while the comparison approach’s performance deteriorates. Gemini thereby enables memory-efficient and informative network integration for large gene networks, and can be used to massively integrate and analyze networks in other domains.

**Availability:** Gemini can be accessed at: https://github.com/MinxZ/Gemini.

## Introduction

Biological networks can provide insights into the underlying mechanisms of human diseases [31, 53, 28, 10, 60, 8, 7, 6, 64, 34, 29], cell differentiation [48, 30, 62], and protein essentiality [68]. As genetic profiling technologies have become more accessible, many large-scale biological networks have been produced, including protein-protein interaction (PPI) networks [30, 55], genetic interaction networks [56], metabolic networks [32, 35], disease networks [2, 67], and patient similarity networks [60]. Importantly, these different types of networks can contain heterogeneous information due to the variation in experimental technology and the underlying biology they measure, although all of the networks capture some components of the complex biological system. Furthermore, each individual network is noisy and incomplete, due to imperfect measurement capabilities. Therefore, there is a pressing need to integrate many biological networks together, thereby pooling information from the many input networks to de-noise the data and combine the distinct sources of information from different experimental measurement types.

Gene networks, where each vertex represents a gene, are one common type of biological network. Such gene networks can be derived from many different experimental sources, including genetic interaction [56], co-expression [14], physical interaction [30], and co-localization [46], among many others. These different types of experimental data show different patterns, which can enhance our biological understanding of genes [68, 49]. In addition, gene networks are often very large, containing several thousand genes each. Although network integration is an active and prolific area of research [12, 13, 61, 21, 18, 65, 15, 41, 40, 43, 44, 26, 27, 57, 58, 59, 31, 53, 28, 10, 51, 11, 8, 37, 33, 9, 6, 7, 64, 34, 22, 16], many exiting network integration methods cannot scale to hundreds of genome-scale gene networks. For example, storing the diffusion states [5, 4] of the 895 human gene networks constructed from BioGRID [42] by GeneMANIA [41] requires 2.77 TB of memory. To be applied to these data, network integration methods must therefore be very memory-efficient. Although some methods can be applied to such data, even recent breakthroughs in unsupervised network integration cannot scale to hundreds of large gene networks due to memory constraints [16]. As recent advances in sequencing technologies have further lowered cost and raised throughput [45], it is likely that even more networks will be available in the future. It is therefore critical that gene network integration methods be scalable to increasing numbers of networks and of genes.

Moreover, existing state-of-the-art unsupervised methods for large biological datasets, such as Mashup [12, 61, 13], assume that important biological signal for a given gene is evenly distributed across networks in the dataset. Although this may hold for smaller, expertly-curated network collections, this is unlikely to extend to hundreds of networks and is not the case in many interactome datasets [38, 20]. Intuitively, different types of networks, such as co-expression and co-localization [54], can encode different features of a gene. In cases where a type of biological evidence is more expensive to experimentally measure, this could cause a data collection to have fewer networks derived from that evidence type. This in turn would diminish the gene-level signal from this biological source.

To address these limitations we present Gemini, an unsupervised network integration approach that has a memory-complexity that is constant to the number of networks. Our key idea is to represent the diffusion state of each network as a vector using fourth-order kurtosis pooling. We then weight each network by an approximation of its uniqueness using this vector, which serves as a more memory-efficient surrogate for the full diffusion state representation. We then simulate many new networks by sampling and mixing-up original networks according to the uniqueness-based weights (**Fig. 1**). This simulation process enables users to either obtain more networks for a more robust integration or consider fewer representative networks for a shorter running time.

**Fig. 1.**
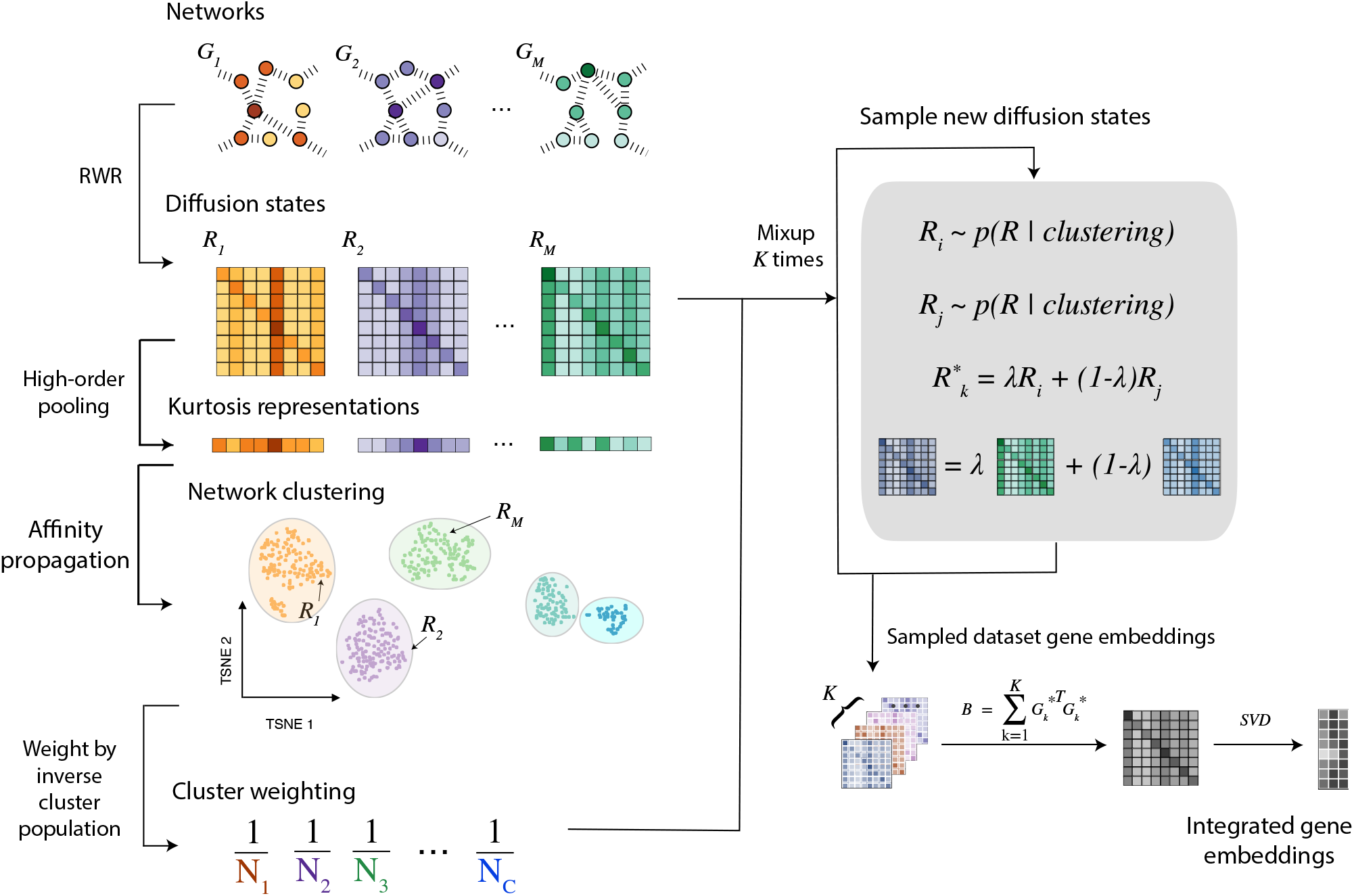
Overview of Gemini algorithm. Given a set of input networks, Gemini uses random walk with restart to compute the diffusion states. Gemini then uses fourth-order kurtosis pooling of the diffusion state matrix as the feature vectors to cluster all networks. Gemini assigns each network a weight inversely proportional to its cluster size. It then randomly samples pairs of networks according to their weights. These pairs of diffusion state matrices are then mixed-up to create a new simulated network collection that more evenly covers the possible diffusion states. We finally aggregate the synthetic dataset and perform an efficient singular value decomposition to produce embeddings for all vertices in the network collection.

We evaluate Gemini on network-based protein function prediction, which has been extensively used to assess biological network integration methods [12, 61, 13, 18, 41, 40, 43, 44, 26, 27, 57, 58, 59, 50, 49, 52]. By integrating hundreds of networks from GeneMANIA [41], we found that Gemini outperforms existing network integration methods by 10% in F_1_ score, 14% in micro-AUPRC, and 71% in macro-AUPRC. Most importantly, we found that the quality of Gemini’s embeddings improved by 27% when additional networks were added to the dataset, while the comparison approach’s [13] performance degraded by 42%. Gemini can therefore balance the many different sources of biological information in the final embeddings while maintaining computational scalability, and can be broadly applied to networks both within and outside of biology for efficient integrative analysis.

## Methods

### Problem definition

We define our input dataset of *M* networks as **𝒢** ^(*M*)^ = *{G*_1_, *G*_2_, …, *G*_*M*_ *}*, where all *G*_*i*_ have the same *n* vertices and can be represented by their adjacency matrix **A**_**i**_. Each network *G*_*i*_ may or may not be weighted or directed. Gemini then outputs an embedding for each vertex, **Z** ∈ ℝ^*n*×*d*^, where *d* is a user-defined embedding dimension, and the embedding 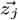 can be used as the feature vector for vertex *j* in downstream tasks.

### Review of Mashup

Gemini’s skeleton is based on the memory-efficient version of Mashup [13, 12], the state-of-the-art network integration approach with memory complexity that is constant in the number of networks and quadratic in the number of nodes, which is critical when integrating many large networks. Fundamentally, Mashup contains three important steps:

1. Compute the diffusion state for each network using random walk with restart (RWR);
2. Integrate the diffusion state for each network;
3. Decompose the integrated states with singular value decomposition (SVD).

In the first step, the diffusion state is calculated from each network’s transition matrix **T**_**i**_ using the RWR algorithm, iteratively computing the state for vertex *j* in network *i* as

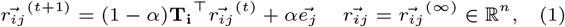

where *α* is a user-specified restart probability and 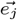 is a standard basis vector, and 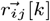 is the probability assigned to vertex *k* given starting state *j* in network *i*. For notational simplicity, we denote the RWR diffusion matrix for all vertices in network *i* as **R**_**i**_ ∈ ℝ^*n*×*n*^.

In the second step, Cho et al. integrate the *M* diffusion state matrices with concatenation, with 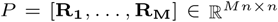. In the third and final step, Cho et al. compute the embedding 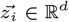 for the *i*th vertex as

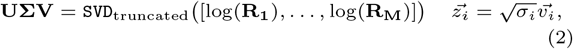

where `SVD`_truncated_ indicates the truncated SVD, and *σ*_*i*_ is the *i*th singular value and *v*_*i*_ is the *i*th right singular vector of the concatenated log-transformed RWR matrices. To reduce memory usage, Cho et al. use the identity that the eigenvectors of **A**^⊤^**A** are the same as the right singular vectors of **A**. Therefore, rather than the *O*(*Mn*^2^) memory requirement for (2), we can instead compute

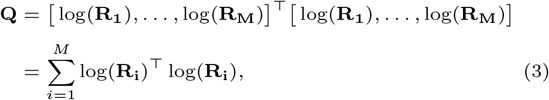

which can be summed one network at a time, thereby only requiring *O*(*n*^2^) memory. Finally, Mashup takes 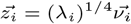, where 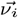 is the *i*th eigenvector of **Q** and *λ*_*i*_ is the corresponding eigenvalue, to efficiently compute the *i*th vertex embedding.

Although this enables fast and scalable integration of large networks, the concatenation of all diffusion state matrices in the second step introduces several important limitations. First, Mashup evenly weights information from all networks during the integration step. When applied to increasingly large number of networks without extensive pre-processing and curation, redundant networks may therefore dominate the downstream embeddings. Second, the networks contained in the input dataset are unlikely to cover the set of possible and reasonable inputs. Therefore, further pooling information from different networks could be useful for further improving downstream vertex embeddings. We aim to address these limitations by refining the second step with Gemini. Gemini maintains the same first step and third step as Mashup.

### Efficient network similarity calculation

Rather than equally combining all networks in the final vertex embedding computation, Gemini weights networks according to their uniqueness, while maintaining efficient memory usage for the large network collection. Mean squared error (MSE) is a widely-used similarity measurement metric. It can therefore be used to quantify the similarity of input networks *G*_*p*_ and *G*_*q*_ based on their diffusion states as

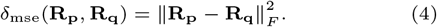

However, in real-world gene network collections there may be too many large networks to run this computation efficiently. Instead, we use RWR to construct a probability distribution centered around the starting vertex, and then quantify the dispersion to each vertex according to its kurtosis, the fourth standardized moment. Compared to low-order pooling (e.g., mean and variance), kurtosis can better capture the tailedness of the diffusion distribution, which can help quantify both global- and local centrality of each vertex in the network, thereby compactly representing its structure. We hypothesize that high-order pooling yields a good approximation for *δ*_mse_(**R**_**p**_, **R**_**q**_) in our real-world datasets, and chose a fourth-order kurtosis poooling. Specifically, we approximate *δ*_mse_(**R**_**p**_, **R**_**q**_) with

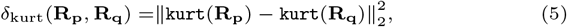

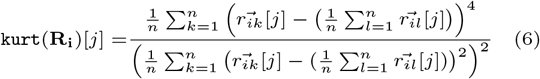

for *j* ∈ *{*1, …, *n}*. Using the kurtosis pooling, we then compute network similarity with *δ*_kurt_(·) in an *O*(*n*) space, rather than an *O*(*n*^2^) space, enabling more network representations to be loaded in memory simultaneously and thereby speeding computation.

### Network weighting according to uniqueness

To identify groups of redundant networks, we then run the affinity propagation clustering algorithm [17] on the kurtosis representations. This does not require a pre-specified number of clusters, but instead computes the number of clusters *C* based on the data. Therefore, if the dataset is already uniformly distributed (all networks are sufficiently dissimilar), they will not be clustered together; however, if the input dataset contains redundancies then similar networks, as defined by (5), will be more likely to cluster together. This step assigns each network to one of the *C* clusters computed by the algorithm as

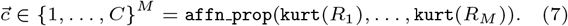

Finally, using *N*_*c*_ to denote the number of input networks assigned to cluster *c*, for all networks *i* ∈ *{*1, …, *M}* we use propensity weighting to define the sampling probability

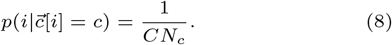

### Diffusion state sampling with mixup

The second limitation that we sought to address is that the input network collections may be unevenly distributed. To address this, Gemini assumes that the diffusion states jointly parameterize a latent understanding of gene similarity. Specifically, Gemini uses the the sampling distribution in (8) as an inverse probability weighting [47]. This leads to instances from different clusters to be sampled equally frequently in expectation. In order to construct a new dataset that can be tuned to the user’s computational resources, we then construct new examples using mixup augmentation [66], simulating

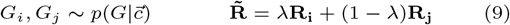

for *λ* ∈ [0, 1] sampled uniformly-at-random. We repeat this process *K* times, where *K* is a hyperparameter defined by the user based on their computational resources, to produce the simulated dataset 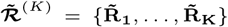, which can better represent possible diffusion state matrices for the input data than the original *M* networks. Although larger values of *K* improve the robustness of 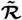, smaller values of *K* enable users with limited computational resources to efficiently integrate input networks with a reduced network collection size.

Using the simulated dataset 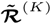, we then repeat memory-efficient embedding step in (3) from Mashup [13] to efficiently compute the vertex representations 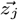 for each vertex *j*. We linearly normalize 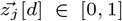 with min-max normalization [19]. As Gemini’s simulated, mixed-up network collection more evenly covers the types of evidence included in the input networks, the vertex embeddings also contain a more even sampling of evidence for all downstream tasks.

### Application to protein function prediction

Biological network integration has been shown to work well in settings where guilt-by-association [1] applies, meaning that connected nodes in the network demonstrate similar phenotypes of interest [39]. One such setting is protein function prediction, which has been used as an important application to validate network integration approaches [12, 61, 13, 18, 41, 40, 43, 44, 26, 27, 57, 58, 59]. We therefore evaluate Gemini’s ability to efficiently generate informative embeddings from a heterogeneous set of input networks using a protein function prediction task, classifying each vertex in a network to a subset of *G* possible functions.

Given the vertex embeddings 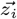, we predict the protein’s function *y*_*i*_ ∈ [0, 1]^*G*^ as

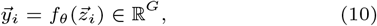

where *σ*(·) represents the sigmoid function. We learn *f*_*θ*_ (·) on a subset of the *n* protein embeddings learned by the unsupervised network integration methods, and evaluate the embedding quality based on the remaining proteins.

## Results

### Dataset, comparison approaches, and experimental settings

We separately consider integration tasks for mouse, human, and yeast datasets to demonstrate Gemini’s applicability to different species with different dataset and network sizes.

#### BioGRID biological networks

We use gene networks for mouse, human, and yeast constructed from BioGRID [42] by Mostafavi et al. [41, 63], with *M* = (403, 895, 505) input networks respectively. Each network has *n* = (21,081, 19,685, 6,365) protein vertices for mouse, human, and yeast respectively.

#### STRING biological networks

We also use gene networks from the STRING dataset [55], where edges indicate an association between two genes based on co-expression data, experimental data, or curated datasets. We use the pre-processed networks from Mashup [13], where network edges are weighted in [0, 1] according to the probability of edge presence. The STRING database yields *M* = 6 input networks each for mouse, human, and yeast, with *n* = (21,310, 17,799, 6,400) protein vertices respectively. Although the STRING database only contains 6 networks for each species, they are curated; in comparison, GeneMANIA’s BioGRID dataset has many more networks for each species, but they are also more noisy.

#### GOA protein function annotations

We used the Gene Ontology Annotation (GOA) database [3, 24] for the downstream protein function prediction task. The GOA dataset comprises *G* = (16,616, 21,656, 8,387) total possible protein function labels for mouse, human, and yeast respectively, and is divided into three sub-ontologies: biological process (BP), molecular function (MF), and cellular component (CC).

#### Mashup and Comparison Models

We compare to the Mashup network integration model [13], the state-of-the-art integration model that can scale to many hundreds of input networks with reasonable computational power. Importantly, we use the memory-efficient version of Mashup that computes integrated gene embedding features with the eigendecomposition in (3), rather than the learnable objective that cannot scale to our datasets. We also compare to *Average Mashup*, where we run the Mashup RWR diffusion and SVD computation in (2) on the average adjacency matrix from the dataset, 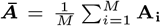. Finally, we also compare to the principal component analysis (PCA) [25] decomposition of ***Ā*** and the truncated SVD of ***Ā***.

In all models, we use *α* = 0.5 for the restart probability in (1), and an embedding dimension *d* = 400 for mouse and human, and *d* = 200 for yeast. During Gemini’s clustering stage, we use Affinity Propagation [17] with a damping factor of 0.875, which we found to converge empirically. For each species’ experiment, we use five-fold cross validation, where 80% of the data is seen during training of each fold. We divide that 80% into a further 80% training and 20% validation set to select the best number of training epochs, use a batch size of 64, and terminate training when the validation loss has not achieved a new global minimum for 50 epochs. We use the same protein function prediction model in (10) for all network integration models, where *f*_*θ*_(·) is implemented as a multi-layer perceptron (MLP) with two hidden layers, with 200- and 100-hidden units respectively, rectified linear unit (ReLU) activation functions, and trained with cross entropy loss.

#### Evaluation Metrics

We evaluate the models’ embedding quality in terms of their test performance on the downstream protein function prediction task. Since this is an imbalanced classification problem, we consider maximum F_1_, which is the F_1_ score with the optimal probability threshold for the model, macro-AUPRC, and micro-AUPRC.

### Gemini improves downstream protein function prediction on BioGRID

We found that Gemini substantially outperformed the comparison approaches [12, 13] on the BP, MF, and CC sub-categories of the GOA annotations for all three species (**Fig. 2**; one-sided paired t-test *p <* 1e−43) by integrating hundreds of networks from BioGRID. Gemini has the largest improvement on human, where it has an average test F_1_ score of 0.50, macro-AUPRC of 0.13, and micro-AUPRC of 0.49, compared to 0.45, 0.07, and 0.43 for the best-performing baseline respectively. The mouse dataset also has a similar improvement, while the yeast improvement is significant but more modest. Since human and mouse networks are larger than yeast networks, this indicates the importance of weighting networks, especially in large network collections. Furthermore, we find that the Average Mashup outperforms Mashup on the BioGRID datasets; we believe that Mashup largely models the more unique networks with the network-specific weight in the SVD computation in (2), rather than sharing the information in the vertex embedding. Gemini addresses this challenge by more evenly sampling different types and combinations of networks, thereby incorporating rare network information into the vertex embeddings as well as the network embeddings.

**Fig. 2.**
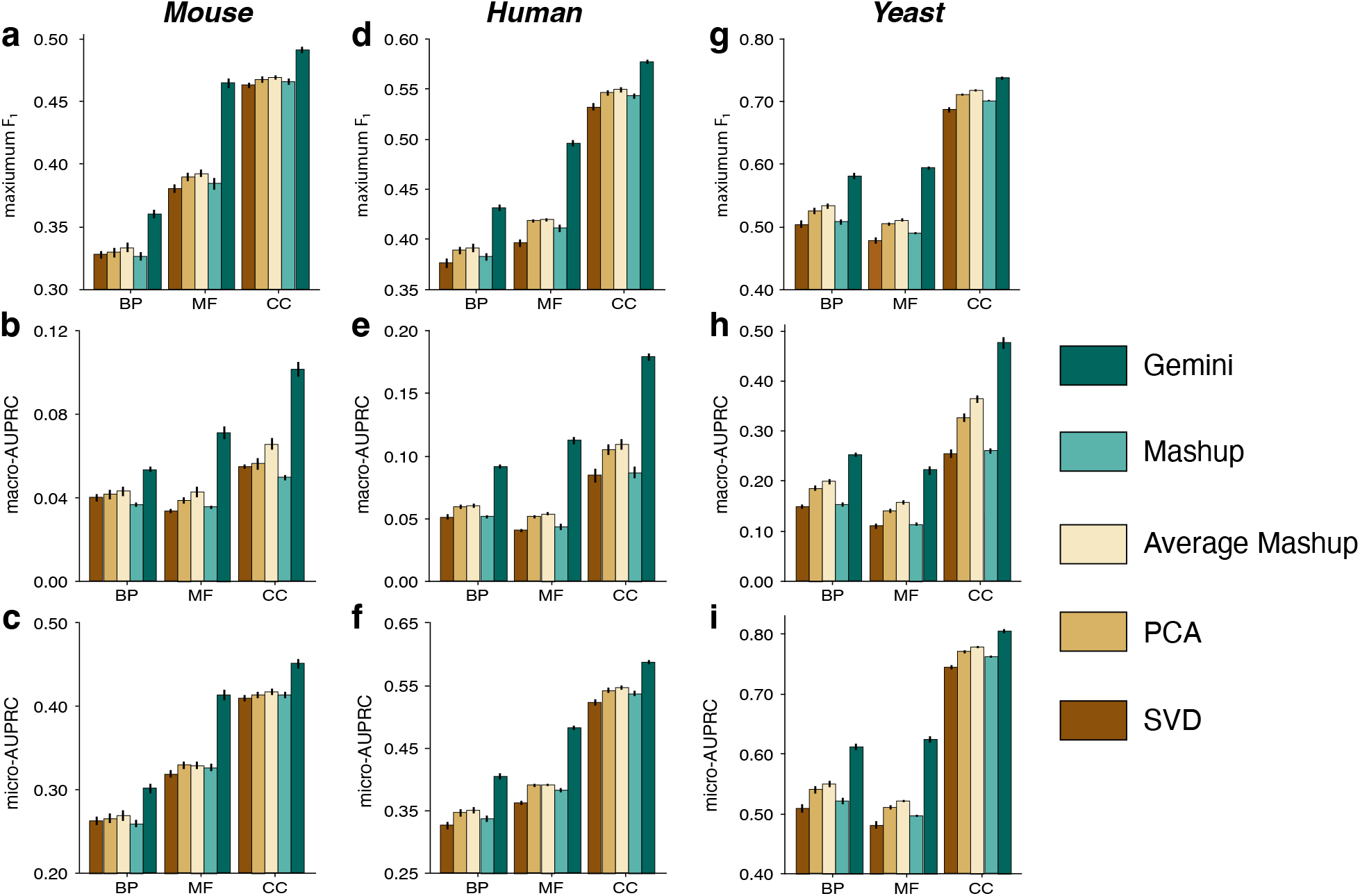
Performance of protein function prediction by integrating networks from BioGRID. **a-i**, Barplot of model performance for protein function prediction task, divided into the BP, MF, and CC sub-ontologies of GOA. Comparison of Mashup, Mashup on ***Ā***, PCA on ***Ā***, SVD on ***Ā***, and Gemini on mouse (**a-c**), human (**d-f**), and yeast (**g-i**) species. We consider F_1_ score (**a**,**d**,**g**), macro-AUPRC (**b**,**e**,**h**), and micro-AUPRC (**c**,**f**,**i**), where higher values indicate better performance and error bars indicate standard error.

### Gemini effectively integrates networks from BioGRID and STRING

We next evaluated Gemini’s performance on integrating all networks from BioGRID and STRING. We repeated the protein function classification task using the GOA database from **Section 3.2**, comparing the quality of the embeddings learned from the STRING network collection, the BioGRID network collection, and the union of the STRING and BioGRID network collections. In such a setting, the union of the two network collections is a superset of the individual collection, and we therefore expect it to contain more information than either collection alone without removing any of the information. We would therefore hope that a network integration method would perform better on the combined network collections than either of the collections alone.

We find that Gemini achieves peak performance on the combined dataset, indicating that it effectively makes use of all networks from two different sources (**Fig. 3**). In particular, Gemini has a maximum F_1_ score of 0.62 on the combined dataset, compared to 0.55 for Mashup. Furthermore, this improves compared to both Gemini’s F_1_ score of 0.58 on the STRING and 0.61 on the BioGRID databases individually. As an important comparison point, we find that Mashup consistently performs best when embeddings are learned only from the STRING networks, and downstream protein function annotation quality degrades when the BioGRID networks are also included. Specifically, while Gemini’s macro-AUPRC performance significantly improves by 27% when BioGRID networks augment the STRING input (two-sided t-test, *p <* 1e−4), Mashup’s performance significantly decreases by 42% (two-sided t-test, *p <* 2e−4). The prominent performance of Gemini when integrating hundreds of networks from many sources demonstrates Gemini’s wide applicability for integrating the accumulating and continually-generated biological networks.

**Fig. 3.**
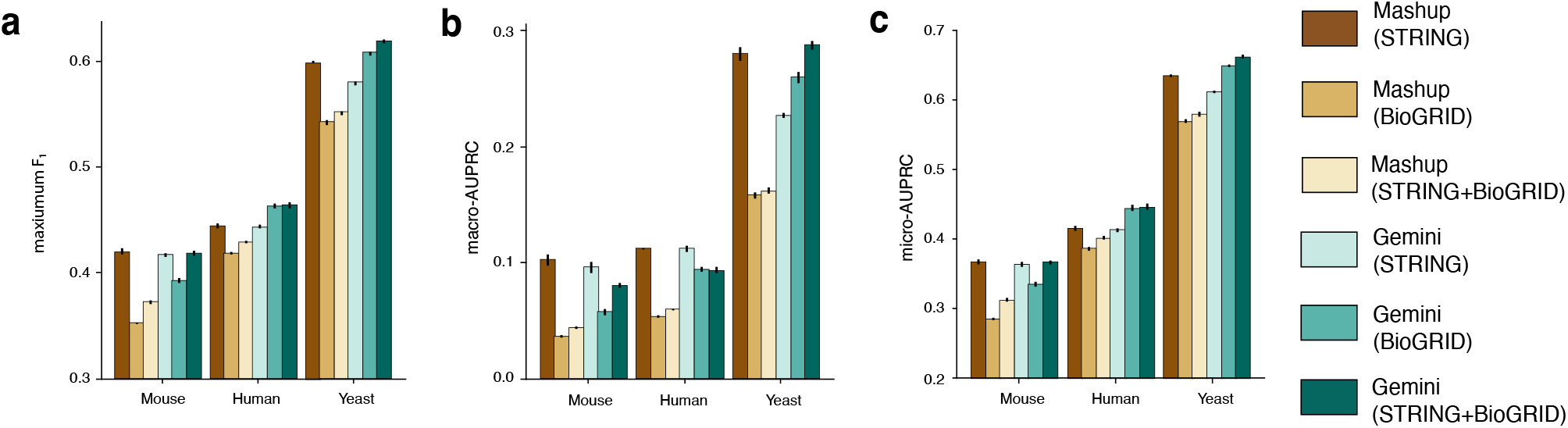
Comparison of network integration for BioGRID and STRING collections. **a-c** Barplots comparing Mashup and Gemini network integration quality on the STRING, BioGRID, and union of STRING and BioGRID network collections, evaluated based on the average performance of the downstream protein function prediction task on GOA. Results for mouse, human, and yeast experiments are shown. Test set performance is measured with the maximum F_1_ score (**a**), macro-AUPRC (**b**), and micro-AUPRC (**c**). Higher values indicate better performance; errors bars indicate the standard error.

### Kurtosis similarity reflects biological similarity

Finally, we sought to further understand the promising performance of Gemini by validating its hypotheses and network representations. We used *δ*_kurt_ in (6) to approximate the slower-to-compute *δ*_mse_ in (4). We empirically observed that kurtosis-based similarity was a good, fast approximation of MSE-based similarity, with a Spearman correlation of 0.87 on the BioGRID yeast dataset (**Fig. 4a**). Gemini is also based on the belief that similar networks reflect biological similarity. We considered the t-Stochastic Neighbor Embedding (t-SNE) [36] of the human BioGRID network’s kurtosis representations, and compared these to the evidence type encoded in the GeneMANIA filenames [41, 63]. We found that the t-SNE space largely clustered according to the type of network (**Fig. 4b**). We also found that the affinity-propagation-based clustering of kurtosis vectors had an adjusted Rand index (ARI) [23] of 0.48 for the human dataset, compared to an ARI of 0 for random cluster assignments, indicating that similar kurtosis representations corresponded to similar biological evidence.

**Fig. 4.**
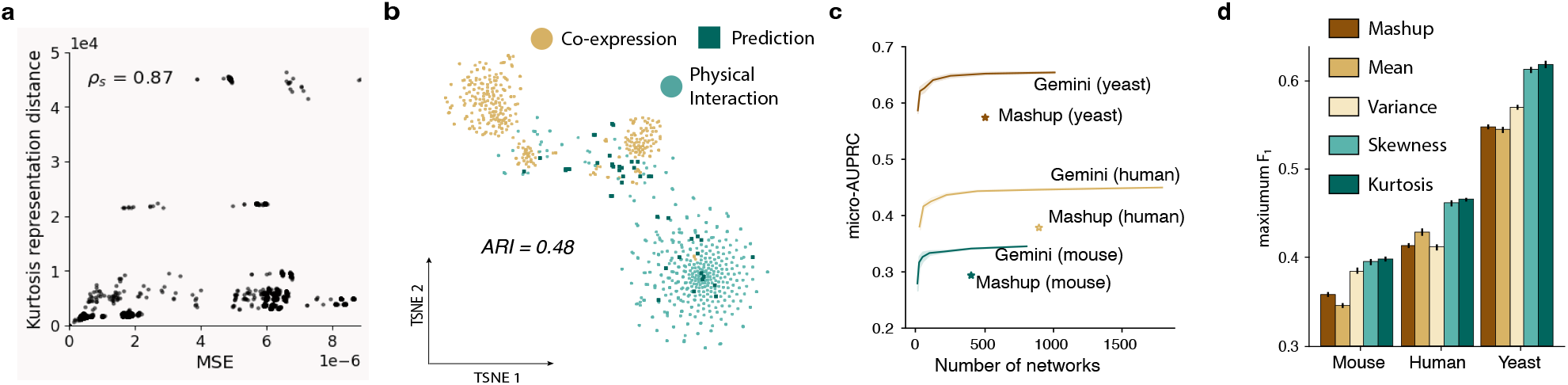
Gemini feature analysis on BioGRID networks. **a**, Scatter plot of MSE- and kurtosis-based similarities for pairs of diffusion state matrices in the yeast dataset, with a Spearman correlation of 0.87. **b**, t-SNE embeddings of kurtosis vector representations for the human dataset, colored by the type of biological evidence used to construct that network, with an adjusted Rand index (ARI) of 0.48. **c**, The average macro-AUPRC (higher is better) with increasing numbers of mixed-up networks *K* for each species across 5 random seeds, compared to Mashup on the original dataset. **d**, Barplot of downstream protein prediction F_1_ score (higher is better) for mean-, variance-, skewness-, and kurtosis-based network similarity approximations, with Mashup included as a reference. Error bars indicate standard error over the test set.

We also evaluated the impact of the size *K* of the synthetic, mixed-up dataset on the protein function prediction performance. We expected that larger dataset sizes would improve downstream performance, but that smaller values of *K* would enable users with smaller computational budgets to compute vertex embeddings. If necessary, we can set *K < M*, reducing the number of networks in the SVD computation compared to the input dataset, and making Gemini run potentially faster than Mashup depending on the computational resources available. We varied *K* from 0.03125 to 2*M* and found that, as expected, larger values of *K* lead to better downstream performance for all species (**Fig. 4c**), although smaller values of *K* can still perform relatively well in resource-limited settings. For instance, the Gemini model with *K* = 0.03125*M* (28 networks) on the human dataset has an F_1_ score 89% of the model trained with *K* = *M*, and this increases to 95% with *K* = 0.0625*M* (56 networks).

Finally, we analyzed the choice of kurtosis pooling by approximating network similarity using different moments. In addition to kurtosis, we compared protein prediction performance based on network similarity approximations using the first moment, with the mean of the diffusion distribution 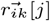; the second central moment, with the variance of 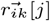; and the third standardized moment, with the skew of 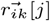. We found that kurtosis and skewness, the fourth and third standardized moments, consistently performed best on all species (**Fig. 4d**), and kurtosis featurization significantly outperforms all other representations (one-sided paired t-test with Bonferroni correction, *p <* 1e−17, 1e−13, 1e−7 for mean, variance, and skewness respectively). This validates our intuition that higher-order pooling can better capture both global and local network structure, compared to low-order pooling measures like mean and variance.

## Conclusion

In this paper we have presented Gemini, a memory-efficient network integration method for large-scale and heterogeneous biological networks. We construct a kurtosis-based pooling of the diffusion state to cover different types and combinations of biological evidence, and use this vector to effectively weight each network. Although we address the under- and over-representation of different types of evidence, Gemini considers cluster membership, and therefore similarity, at the network-level. Future work could extend this notion to a vertex-based similarity, where two networks could be considered similar for some genes and different for others. Furthermore, as Gemini will weight unique networks more heavily than repetitive networks, outlier detection methods could be employed to remove particularly noisy or adversarial networks from consideration.

## Author contributions statement

A.W., M.Z. and S.W. conceived the experiment(s), A.W. and M.Z. conducted the experiment(s), A.W. and S.W. wrote the manuscript with feedbacks from other authors.

